# A Neural Signature of Regularity in Sound is Reduced in Older Adults

**DOI:** 10.1101/2021.02.18.431898

**Authors:** Björn Herrmann, Burkhard Maess, Ingrid S. Johnsrude

## Abstract

Sensitivity to repetitions in sound amplitude and frequency is crucial for sound perception. As with other aspects of sound processing, sensitivity to such patterns may change with age, and may help explain some age-related changes in hearing such as segregating speech from background sound. We recorded magnetoencephalography to characterize differences in the processing of sound patterns between younger and older adults. We presented tone sequences that either contained a pattern (made of a repeated set of tones) or did not contain a pattern. We show that auditory cortex in older, compared to younger, adults is hyperresponsive to sound onsets, but that sustained neural activity in auditory cortex, indexing the processing of a sound pattern, is reduced. Hence, the sensitivity of neural populations in auditory cortex fundamentally differs between younger and older individuals, overresponding to sound onsets, while underresponding to patterns in sounds. This may help to explain some age-related changes in hearing such as increased sensitivity to distracting sounds and difficulties tracking speech in the presence of other sound.

## Introduction

Many adults aged 50 or older experience challenges understanding speech in the presence of background sound (Pichora-Fuller, 2003; Pichora-Fuller et al., 2016), but the underlying neural sources contributing to such deficits are not fully understood. Speech contains rich, regular patterns, such as quasi-regular amplitude fluctuations at 4–5 Hz (Rosen, 1992; Varnet et al., 2017), and perceptual sensitivity to sound pattern and speech-in-noise perception correlate with each other (Holmes and Griffiths, 2019), suggesting shared mechanisms (Holmes et al., 2021). The perceptual processes through which sensitivity to such patterns may contribute to speech perception likely include the segregation of unique, concurrent sound streams (Schröger, 2005, 2007; Snyder and Alain, 2007; Winkler et al., 2009; Bendixen, 2014) and the recognition and prediction of relevant sound features (Jones and Boltz, 1989; Nobre et al., 2007; Henry and Herrmann, 2014; Nobre and van Ede, 2018). The current study is concerned with the degree to which patterns are represented in the brains of older individuals and whether neural sensitivity to patterns differs between younger and older adults.

The detection of a regular pattern in a sound is associated with an increase in a sustained, low-frequency, DC power offset in cortical electroencephalography (EEG) and magnetoencephalography (MEG) recordings (Barascud et al., 2016; Southwell et al., 2017; Herrmann and Johnsrude, 2018b). Sustained neural activity manifests as soon as a pattern, such as repetition of a set of tones, is present (Southwell et al., 2017; Herrmann and Johnsrude, 2018b; Southwell and Chait, 2018; Herrmann et al., 2021). It also manifests for spectrally coherent chord fluctuations (Teki et al., 2016), complex sounds made of isochronous tone sequences (Sohoglu and Chait, 2016), and repeated amplitude or frequency modulations (Gutschalk et al., 2002; Ross et al., 2002; Herrmann and Johnsrude, 2018b; Herrmann et al., 2019). Sustained activity increases with the degree of regularity of a pattern, for example, with increasingly coherent frequency modulation in sounds (Teki et al., 2016; Herrmann and Johnsrude, 2018b). The magnitude of sustained activity is thought to reflect prediction-related processes (Heilbron and Chait, 2018).

Accumulating evidence suggests that aging and age-related hearing loss are associated with a loss of inhibition throughout the auditory pathway following peripheral decline (Caspary et al., 2008; Rabang et al., 2012; Ouellet and de Villers-Sidani, 2014). This may render neurons in the aged auditory system hyperresponsive to sound (Hughes et al., 2010; Alain et al., 2012; Bidelman et al., 2014; Overton and Recanzone, 2016; Presacco et al., 2016b, a; Herrmann et al., 2018) and shorten the time it takes for neurons to regain responsiveness following adaptation to sound (de Villers-Sidani et al., 2010; Mishra et al., 2014; Herrmann et al., 2016; Herrmann et al., 2019). Changes in inhibition, responsivity, and adaptation associated with aging and hearing loss likely affect all aspects of hearing (Herrmann and Butler, 2021), including sensitivity to sound patterns.

Some initial evidence suggests that sustained neural activity may be reduced in older compared to younger people. Many years ago, Pfefferbaum and colleagues (1979) demonstrated that sustained activity elicited by a short sine tone is reduced for older compared to younger adults. More recent work indicates that younger individuals exhibit pattern-related sustained activity in response to amplitude-modulated sounds, whereas older adults do not appear to, although the difference between these groups was not significant (Herrmann et al., 2019). Another study yielded data suggestive of reduced sustained activity in older compared to younger people in response to repeated tone sequences (Al Jaja et al., 2020), but stimulus parameters differed between age groups in this paper. A controlled experiment with sufficient power is thus required to elucidate whether sustained neural activity to regular sound patterns differs between younger and older people.

Previous work investigating sustained neural activity in older adults has utilized low-density electroencephalography (EEG; fewer than 20 electrodes; Pfefferbaum et al., 1979; Herrmann et al., 2019; Al Jaja et al., 2020). This type of EEG is not very well suited for the localization of neural sources generating scalp-recorded signals. Magnetoencephalography typically allows for better source reconstruction than EEG, because magnetic fields are less distorted by the skull and scalp than the EEG-recorded electric potentials (Hämäläinen et al., 1993; Hämäläinen and Hari, 2002). Previous MEG source localizations in younger adults suggest that the auditory cortex underlies sustained neural activity (Hari et al., 1980; Pantev et al., 1994; Pantev et al., 1996; Gutschalk et al., 2002; Ross et al., 2002; Okamoto et al., 2011; Barascud et al., 2016; Teki et al., 2016) and that additional brain regions in parietal cortex, frontal cortex, and hippocampus may also contribute (Tiitinen et al., 2012; Barascud et al., 2016; Teki et al., 2016). Whether the neural sources of pattern-related sustained activity differ between younger and older adults is unknown.

In the current study we recorded MEG from younger and older adults while they listened to sound sequences. Sequences were made by taking pure tones at different frequencies and either repeating the same small set of these in the same order, so that a regular pattern is heard, or by presenting them pseudo-randomly so that no pattern is present. We investigate whether sustained neural activity to a regular sound pattern differs between younger and older individuals. We also examine whether auditory cortex is generally more responsive to sound in older, compared to younger adults, as has been previously reported (Bidelman et al., 2014; Herrmann et al., 2018).

## Methods and Materials

### Participants

Twenty-six younger (mean: 26.7 years; range: 21–33 years; 13 males and 13 females) and twenty-five older adults (mean: 63.9 years; range: 53–73 years; 11 males and 14 females) participated in the current study. Participants reported no neurological disease or hearing impairment, gave written informed consent, and were paid for their participation. None of the participants wore a hearing aid or reported having been prescribed a hearing aid. We focused on a typical sample of older individuals, allowing for the possibility of some degree of hearing impairment. The study was conducted in two sessions on separate days (range: 1-43 days apart; median: 7 days apart; no age-group difference: t_49_ = 0.99, p = 0.327). The study was conducted in accordance with the Declaration of Helsinki, the Canadian Tri-Council Policy Statement on Ethical Conduct for Research Involving Humans (TCPS2-2014), and was approved by the local Nonmedical Research Ethics Board of the University of Western Ontario (protocol ID: 106570).

### Hearing assessment and hearing thresholds

Pure-tone audiometric data were acquired for each participant (Figure 1). The pure-tone average hearing threshold (i.e., the mean across the 0.25, 0.5, 1, 2, and 4 kHz frequencies) was larger for older compared to younger adults (t_49_ = 7.79, p = 4×10^−10^, r_e_ = 0.744; Figure 1, right). This indicates a mild-to-moderate hearing impairment in many of the older adults and is consistent with the high-frequency sloping loss characteristic of age-related hearing impairment (Moore, 2007; Plack, 2014) as well as with previous electrophysiological studies that investigated differences in sound processing between younger and older adults (Presacco et al., 2016b; Herrmann et al., 2018).

**Figure 1:**
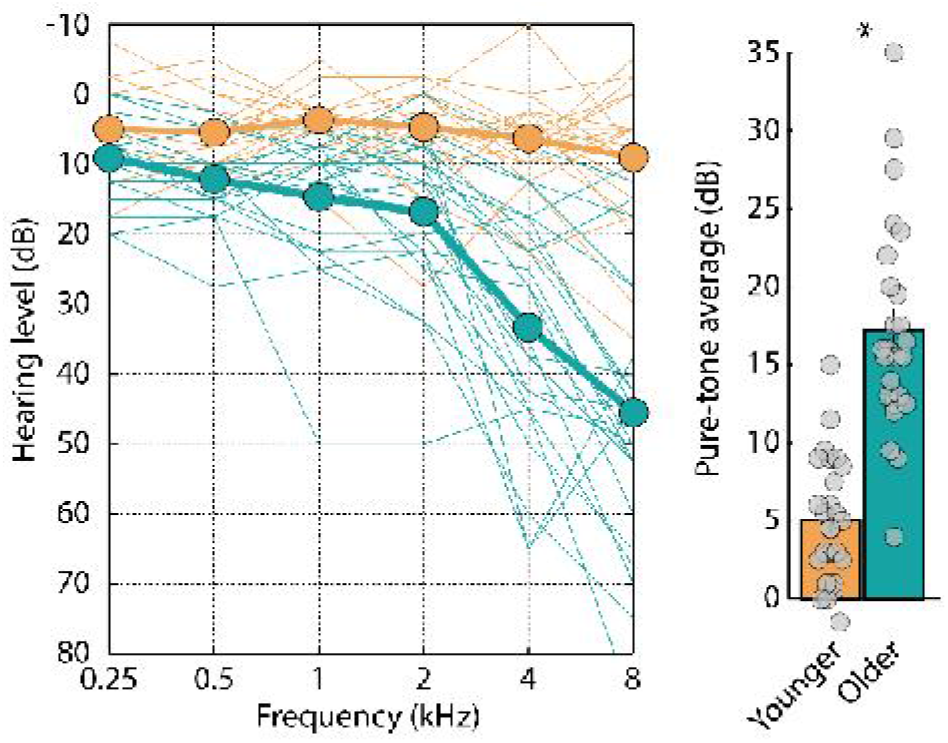
Audiograms and pure-tone average hearing threshold. Left: Audiograms for each participant. Thin lines reflect individual participant data. Thick lines reflect the mean across participants. Right: Pure-tone average hearing threshold (mean across 0.25, 0.5, 1, 2, and 4 kHz). Gray dots reflect the threshold for individuals.

For each participant, we measured the hearing threshold (i.e., sensation level [SL]) using a method-of-limits procedure (Herrmann and Johnsrude, 2018a; Herrmann et al., 2019) as a reference threshold in MATLAB software for sound presentation. Participants listened to a 12-s pure tone with a frequency of 1323 Hz that changed continuously in intensity at a rate of 5 dB/s (either decreased [i.e., starting at suprathreshold levels] or increased [i.e., starting at subthreshold levels]). Participants pressed a button when they could no longer hear the tone (intensity decrease) or when they started to hear the tone (intensity increase); the sound stopped after button press. The sound intensity at the time of the button press was noted for 6 decreasing sounds and 6 increasing sounds (decreasing and increasing sounds alternated), and these were averaged to determine the individual hearing threshold. The mean hearing threshold was elevated for older compared to younger adults (t_49_ = 5.208, p = 3.7×10^−6^, r_e_ = 0.597), which was expected given the audiograms (Figure 1).

All acoustic stimuli described below were presented at 55 dB above each individual’s hearing threshold – that is, at 55 dB sensation level – in order to control for audibility across age groups. Because hearing thresholds were on average elevated for older compared to younger adults, sounds during the MEG recordings were on average more intense in sound-pressure level (SPL) in older compared to younger individuals. Higher sound levels can lead to larger brain responses to sound onsets (Picton et al., 1974; Picton et al., 1978; Pfefferbaum et al., 1979; Polich et al., 1988; Schadow et al., 2007; Herrmann et al., 2018) as well as for sustained activity (Picton et al., 1978; Pfefferbaum et al., 1979). Because we hypothesized that regularity-related sustained activity would be smaller for older compared to younger adults, playing sounds at a higher level for older adults only works against this hypothesis. Presenting sounds at sensation level was thus favorable in the current study. However, a higher sound level for older compared to younger adults could bias statistical analyses for investigations of age-related hyperresponsivity to sound, for which we expect larger responses in older compared to younger adults. Hence, for these analyses, we also used a subgroup of 14 participants of each age group for which the hearing threshold – and thus the sound level of the acoustic presentation – did not differ (t_26_ = 0.956, p = 0.348, r_e_ = 0.184; younger mean [±sd]: −94.16 dB ±1.39, older mean [±sd]: −93.36 dB ±2.65^1^) to confirm our results.

### Acoustic stimulation and procedure

Acoustic stimuli were 4-s long sequences that each consisted of 96 pure-tone pips arranged in twelve sets of eight tones each (see also Barascud et al., 2016; Herrmann and Johnsrude, 2018b; Southwell and Chait, 2018; Herrmann et al., 2021). Each set had a duration of 0.333 s. Pips were 0.0417 s in duration with attack and decay times of 0.007 s, and no gap between tones, or sets. The frequency of each tone was one of 150 possible values between 700 and 2500 Hz (logarithmically spaced).

Acoustic stimuli were presented in two conditions, ‘Pattern-Absent’ and ‘Pattern-Present’, which occurred with equal probability (50%). In the ‘Pattern-Absent’ condition, tones with different frequencies were presented in pseudo-random order without a pattern, whereas in the ‘Pattern-Present’ condition, tones transitioned from random to a regular pattern 1 s (3 sets) after sound onset. For the ‘Pattern-Absent’ condition, 8 new frequency values were randomly selected for each of the 12 sets (Figure 2, top). In the ‘Pattern-Present’ condition, 8 new frequency values were randomly selected for each of the first 3 sets (0–1 s; similar to ‘Pattern-Absent’), and then 8 new random frequency values were selected and repeated in the same order for the remaining 9 sets, thereby creating a regular pattern (Figure 2, bottom). These conditions are similar to the sounds used in previous studies that investigated sustained neural activity (Barascud et al., 2016; Southwell et al., 2017; Herrmann and Johnsrude, 2018b).

**Figure 2:**
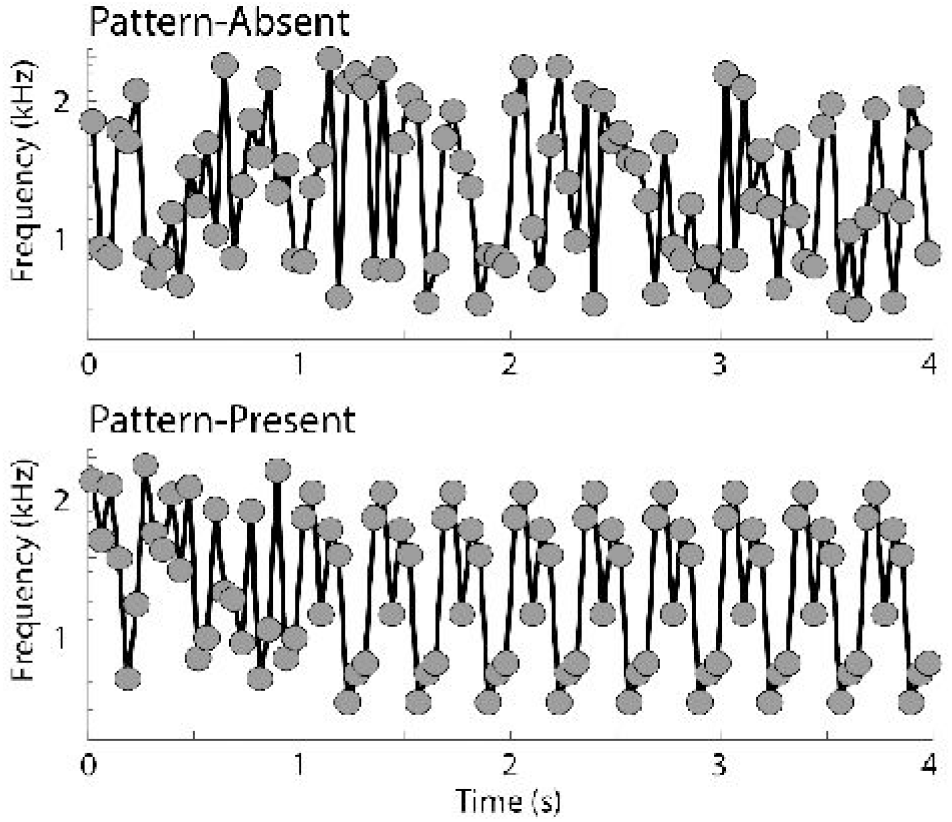
Schematic of acoustic stimulation for ‘Pattern-Absent’ and ‘Pattern-Present’ conditions. Sound frequency is displayed on the y-axis and dots reflect the sound frequency of individual tones of the tone sequence.

In each of the two recording sessions, participants were presented with one 12-min block of stimulation as part of recording sessions for an additional project not presented here. The data from the experimental blocks reported here were recorded in the beginning of the recording sessions. Participants listened passively to sixty 4-s sound sequences of each condition per session, while watching a muted movie of their choice, with subtitles, that was projected into the electromagnetically shielded room via a mirror system. Trials of the Pattern-Absent and the Pattern-Present conditions were presented pseudo-randomly throughout the block, such that each condition could occur maximally three times in direct succession. Across both sessions, participants listened to 120 trials per condition. Trials were separated by a 2-s inter-stimulus interval.

### Magnetoencephalographic recordings and initial preprocessing

Magnetoencephalographic data were recorded using a 306-channel Neuromag Vectorview MEG (MEGIN Oy, Helsinki, Finland; sampling rate: 1000 Hz, online filter: DC–330 Hz) at the Max Planck Institute for Human Cognitive and Brain Sciences in Leipzig, Germany. Data were recorded in an electromagnetically shielded room (AK3b, Vacuumschmelze, Hanau, Germany). The signal space separation (SSS) method (maxfilter© version 2.2.15; default parameter setting L_in_ = 8; L_out_ = 3) was used to suppress external interference, interpolate bad channels, and transform each person’s individual data to the sensor space of the first block of the first session to ensure the data are in a common space (Taulu et al., 2004; Taulu et al., 2005).

### Combination of magnetometer and gradiometer channels

The Vectorview MEG device records magnetic fields using 102 magnetometers and 204 gradiometers in 102 locations distributed around the head. In order to account for all data that were recorded, we combined signals from magnetometer and gradiometer channels (Herrmann et al., 2018). Magnetometers and gradiometer differ in their configuration, such that magnetometers measure magnetic fields in Tesla (T), while gradiometers (a coupled pair of magnetometers) measure differences in the same magnetic fields over a distance of 0.0168 m in Tesla per meter (T/m). The combination of channel types requires accounting for their different units. We transformed all channels into magnetometer channels, because such a model only requires a linear interpolation that results in the same unit for all channels. To this end, we applied the following transformation matrix to each of the 102 sensor triplets (i.e., one triplet comprises two gradiometer channels and one magnetometer channel):

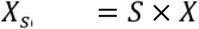

where X consists of a 3 × n matrix (with n being the number of data samples over time). The three rows of X refer to the two gradiometers and one magnetometer (i.e., one triplet). S refers to a 5 × 3 scaling matrix with the following elements:

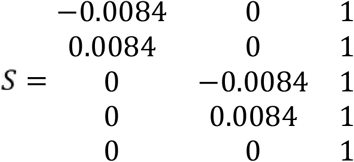

The value 0.0084 reflects half of the distance between the two gradiometer loops measured in meters, and the transformation constitutes a linear approximation of the magnetic field at each of the triplets. The transformation replaces the sensor triplet by a sensor quintet of magnetometers. The columns of S refer to the triplet of two gradiometers and one magnetometer and the rows of S refer to the resulting five magnetometers. This procedure resulted in signals from 510 magnetometer channels centered on and around 102 locations around a participant’s head (Herrmann et al., 2018).

### Preprocessing of magnetoencephalographic data

Data were high-pass filtered (0.7 Hz; 2391 points, Hann window), low-pass filtered (20.3 Hz, 119 points, Kaiser window), down-sampled to 250 Hz, and divided into 6-s long epochs time-locked to sound onset (from 1 s before to 5 s after sound onset). Independent components analysis (runica method, Makeig et al., 1996; logistic infomax algorithm, Bell and Sejnowski, 1995; Fieldtrip implementation, v20130727, Oostenveld et al., 2011) was used to identify and remove activity related to blinks, horizontal eye movements, muscle activity, and noisy channels. Identification of components related to these non-brain activities was done manually through visual inspection of component time courses, topographies, and frequency spectra by BH. Epochs in which a signal change larger than 8 Picotesla (pT) occurred in any channel were excluded. The remaining data were used to investigate age differences in evoked responses to the onset of the sounds.

In order to investigate the sustained neural activity, the same pipeline was computed a second time, with the exception that high-pass filtering was omitted. Omission of the high-pass filter is necessary to investigate sustained activity, because the response is a very low-frequency signal reflecting a DC shift (Barascud et al., 2016; Southwell et al., 2017; Herrmann and Johnsrude, 2018b). Activity related to blinks, horizontal eye movements, muscle activity, and noisy channels was removed using the identified components from the high-pass filtered data. Epochs in which a signal change larger than 8 pT occurred in any channel were excluded.

### Analysis of responses to sound onset

High-pass filtered data were used to investigate whether neural responses to the onset of the sounds differed between age groups. This analysis aimed to test whether the auditory cortex of older adults is hyperresponsive to sound, consistent with reduced inhibition (Caspary et al., 2008; Hughes et al., 2010; Juarez-Salinas et al., 2010). Data from the Pattern-Absent and Pattern-Present conditions were averaged because both conditions were identical for the first second of the sound. Epochs ranging from −0.15 s to 0.5 s time-locked to sound onset were extracted. Absolute values were calculated for signals of each channel (because magnetic fields have opposite polarities in directions perpendicular to the tangential orientation aspect of the underlying neural source). The mean signal from the pre-stimulus period (−0.15 to 0 s) was subtracted from the signal at each time point, separately for each channel (baseline correction). Responses were averaged across channels, resulting in one response time course per participant.

For the statistical analysis, differences in response amplitude between age groups were assessed for each time point using independent samples t-tests. False discovery rate was used to account for multiple comparisons (Benjamini and Hochberg, 1995; Genovese et al., 2002). We confirmed the results with two independent samples t-tests that contrasted the amplitudes of the M50 (0.03–0.06 s) and M100 (0.09–0.13 s) between age groups, which have previously been shown to differ between younger and older adults (Sörös et al., 2009; Alain et al., 2012; Herrmann et al., 2018).

### Analysis of pattern-related sustained activity

Non-high-pass filtered data were used to investigate whether sustained neural activity associated with a pattern in sounds differs between age groups. The 6-s epochs (−1 to 5 s, time-locked to sound onset) were used. Absolute values were calculated for signals of each channel and the mean signal from the pre-stimulus period (−1 to 0 s) was subtracted from the signal at each time point, separately for each channel (baseline correction). Responses were averaged across channels, resulting in one response time course per condition and per participant.

Statistical analysis focused on responses during the last half of each stimulus: the 2–4 s time window. By 2 s, the repeating set of tones would have been presented 3 times (2 full repetitions) in the Pattern-Present condition (Barascud et al., 2016; Teki et al., 2016; Herrmann and Johnsrude, 2018b). An ANOVA with the within-subjects factor Condition (Pattern-Absent, Pattern-Present) and the between-subjects factor Age Group (younger, older) was calculated.

### Source localization of magnetoencephalographic data

Anatomically constrained source localization was used to localize the sources underlying the neural activity in sensor space. Individual T1-weighted MR images (3T Magnetom Trio, Siemens AG, Germany) were available for each participant. The MR images were used to construct inner skull surfaces (volume conductor) and mid-gray matter cortical surfaces (source model; using Freesurfer and MNE software; https://surfer.nmr.mgh.harvard.edu/; http://www.martinos.org/mne/). The MR and the MEG coordinate systems were co-registered using MNE software, which included an automated and iterative procedure that fitted the >300 digitized head surface points (Polhemus FASTRAK 3D digitizer) to the MR reconstructed head surface (Besl and McKay, 1992). The inner skull was extracted from the MR images using MNE software and used to calculated lead fields using the boundary element model as implemented in Fieldtrip software (Nolte, 2003). Inverse solutions were calculated using the sLORETA method (Pascual-Marqui, 2002). Neural activity was spatially smoothed across the surface using an approximation to a 6-mm FWHM Gaussian kernel (Han et al., 2006). Individual cortical representations were transformed to a common coordinate system (fsaverage standard brain; Fischl et al., 1999b). Workbench software (v1.4.2; https://www.humanconnectome.org/) was used for visualization of source localizations morphed to the pial cortical surface of the fsaverage standard brain (Fischl et al., 1999a). Source localizations were calculated for onset responses and for sustained neural activity. In order to visualize and analyze pattern-related auditory cortex activity, we averaged source-localization amplitudes across regions of the superior temporal plane (A1, A4, PBelt, MBelt, and LBelt) using the brain parcelations of the Human Connectome Project (Glasser et al., 2016).

### Effect sizes

Effect sizes are provided as partial η^2^ for ANOVAs and as r_e_ (r_equivalent_) for t-tests (Rosenthal and Rubin, 2003). r_e_ is equivalent to the square root of partial η^2^ for ANOVAs.

### Data availability

This study was not pre-registered. MEG data in BIDS format (Pernet et al., 2019) are available at https://figshare.com/projects/A_Neural_Signature_of_Regularity_in_Sound_is_Reduced_in_Older_Adults/121803.

## Results

### Responses to sound onset are enhanced in older compared to younger adults

Figure 3A displays the neural response time courses elicited by the onset of the sounds. Responses were larger in older compared to younger adults in the M50 and M100 time windows (black line in Figure 3A, FDR-thresholded). Figure 3B/C shows the mean amplitudes and topographical distributions for the M50 and M100 time windows. Larger neural responses for older compared to younger adults were also observed for the subgroups of 14 participants per age group for which hearing thresholds – and thus sound-presentation levels – did not differ (M50: t_26_ = 4.812, p = 5.5×10^−5^, r_e_ = 0.686; M100: t_26_ = 4.257, p = 2.3×10^−4^, r_e_ = 0.641; all participants: M50: t_49_ = 6.295, p = 8.2×10^−8^, r_e_ = 0.669; M100: t_49_ = 4.015, p = 2×10^−4^, r_e_ = 0.497). Finally, regressions calculated to predict M50 or M100 responses from age group, while including sensation-level threshold and audiometric pure-tone average as co-variates, also revealed an effect of age group (M50: t_47_ = 3.199, p = 0.002; M100: t_47_ = 2.571, p = 0.013). These results demonstrate that even when sound level does not differ between younger and older adults, older adults exhibit hyperresponsiveness to sound. There was no age difference for the M200 (0.16–0.22 s: t_49_ = 1.088, p = 0.282). Source localizations show activity in superior temporal cortex, including auditory cortex, underlying M50 and M100 responses in both age groups (Figure 3D/E).

**Figure 3:**
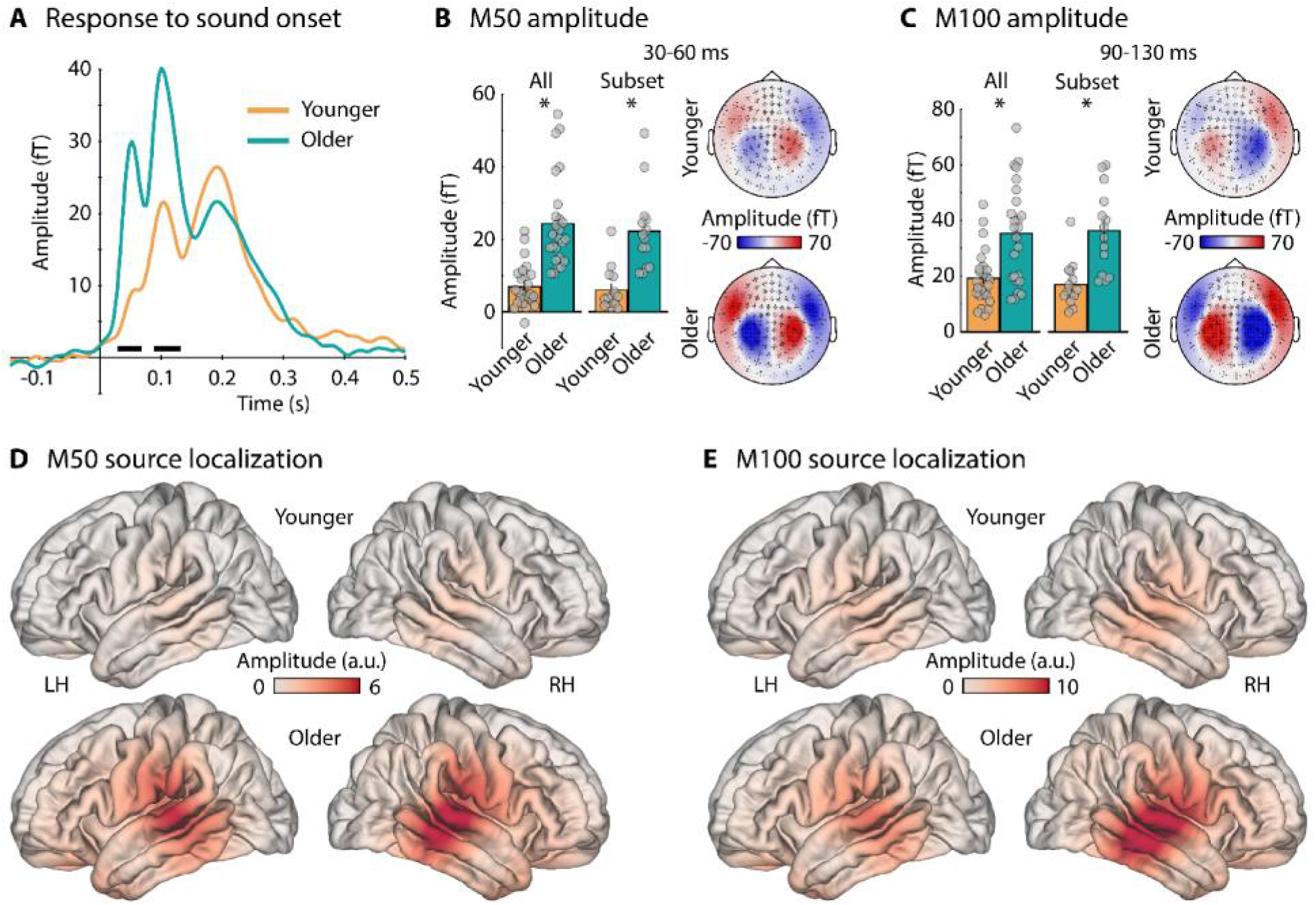
Neural responses to the onset of sounds. A: Time courses of neural activity (root-mean square amplitude, averaged across all channels). The black line indicates a significant difference between age groups (FDR-thresholded). B: Mean activity and topographies for the M50 time window (30–60 ms) for all participants and the subset of 14 participants for which sound level did not differ between younger and older adults. C: Mean activity and topographies for the M50 time window (90–130 ms) for all participants and the subset of 14 participants. D: Source localization for the M50 time window. E: Source localization for the M100 time window. *p ≤ 0.05

### Pattern-related sustained activity is reduced in older compared to younger adults

Figure 4A and B show response time courses and topographical distributions for the Pattern-Absent and the Pattern-Present condition for both age groups. The ANOVA for the 2-4 s time window revealed a Condition × Age Group interaction (F_1,49_ = 9.839, p = 0.003, n_p_^2^ = 0.167; also significant for the subset of participants for which sound level did not differ: F_1,26_ = 6.792, p = 0.015, n_p_^2^ = 0.207): While both age groups show larger sustained activity for the Pattern-Present compared to the Pattern-Absent condition (younger: F_1,25_ = 49.692, p ≤ 1×10^−6^, n_p_^2^ = 0.665; older: F_1,24_ = 6.287, p = 0.019, n_p_^2^ = 0.208), this difference was larger in younger compared to older adults (Figure 4C). There was no difference between age groups for the Pattern-Absent condition (F_1,49_ = 0.528, p = 0.471, n_p_^2^ = 0.011). The main effect of Condition (F_1,49_ = 45.185, p ≤ 1×10^−6^, n_p_^2^ = 0.48) and the main effect of Age Group (F_1,49_ = 6.994, p = 0.011, n_p_^2^ = 0.125) were also significant.

**Figure 4:**
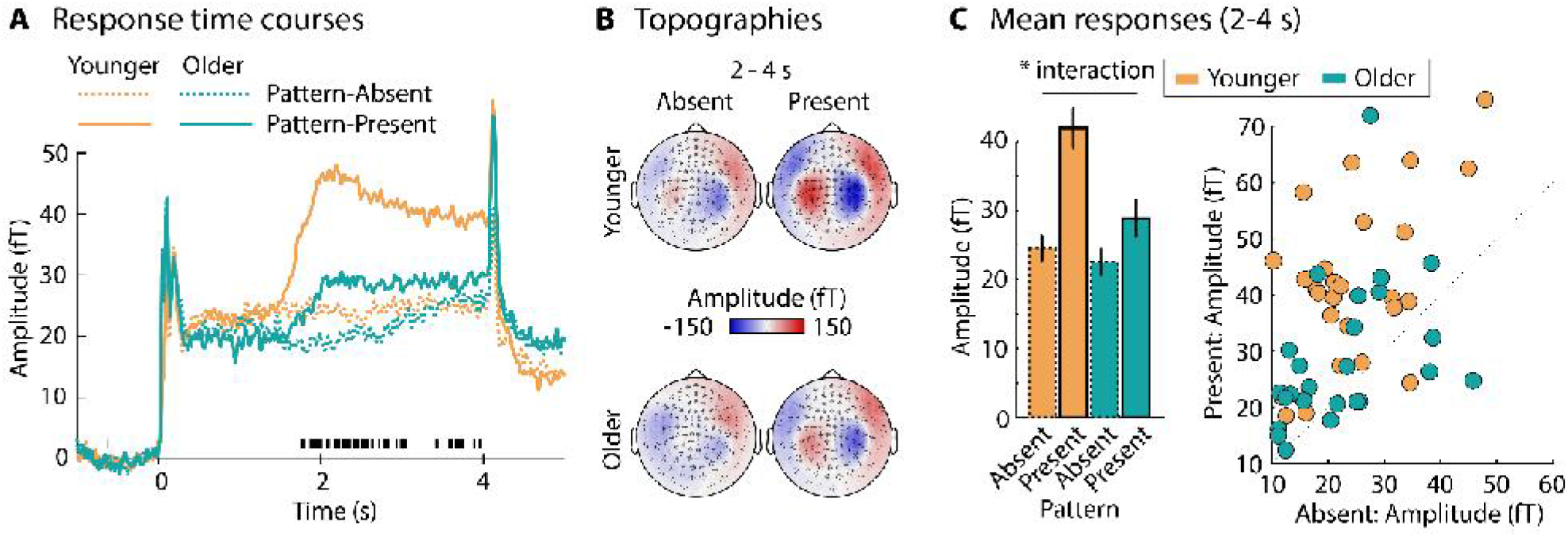
Pattern-related sustained activity. A: Response time courses (root-mean square amplitude, averaged across all channels). The black markings below the time courses indicate the time points at which the Condition × Age Group interaction was significant (FDR-thresholded). B: Topographical distributions for each condition and age group for the 2-4 s time window. C: Mean responses in the 2-4 s time window. Bar graphs reflect the mean across participants. Error bars are the standard error of the mean. Data points for each participant are shown on the right. Data points above the diagonal (dashed line) reflect a larger response for the Pattern-Present compared to the Pattern-Absent condition.

In order to explore the relation between the response to sound onset and the regularity-related sustained activity effects in older adults, we calculated the difference between the Pattern-Present and the Pattern-Absent conditions and correlated the response difference with the M50 and M100 responses to sound onset. Correlations were not significant (M50: r = −0.075, p = 0.722, df = 23; M100: r = −0.011, p = 0.957, df = 23). However, the relation between hyperactivity in response to sound and hearing loss is non-linear (Qiu et al., 2000; Salvi et al., 2017; Herrmann and Butler, 2021), and we may thus not expect a linear correlation between hyperactivity and changes in regularity-related sustained activity in older adults.

Source localizations revealed that the strongest activity associated with pattern-related sustained activity was present in superior temporal cortex and auditory cortex (Figure 5A). Indeed, we observed the same interaction for auditory cortex activity (F_1,49_ = 10.68, p = 0.002, n_p_^2^ = 0.179; Figure 5B/C; for the subset of participants: F_1,26_ = 7.299, p = 0.012, n_p_^2^ = 0.219) that we observed in sensor space (Figure 4C), such that the increase in sustained activity for the Pattern-Present compared to the Pattern-Absent condition was significant for both age groups (younger: F_1,25_ = 50.652, p ≤ 1×10^−6^, n_p_^2^ = 0.670; older: F_1,24_ = 23.833, p = 5.6×10^−5^, n_p_^2^ = 0.498), with a larger difference in younger compared to older adults. In contrast to the sensor space data of sustained activity, sustained activity in auditory cortex elicited by the Pattern-Absent condition was also larger for younger compared to older adults (F_1,49_ = 4.704, p = 0.035, n_p_^2^ = 0.088; Figure 5B/C), consistent with observations of reduced sustained activity to a sine tone in older compared to younger adults (Pfefferbaum et al., 1979). Main effects of Condition (F_1,49_ = 73.205, p ≤ 1×10^−6^, n_p_^2^ = 0.599) and Age Group (F_1,49_ = 10.176, p = 0.002, n_p_^2^ = 0.172) were also significant.

**Figure 5:**
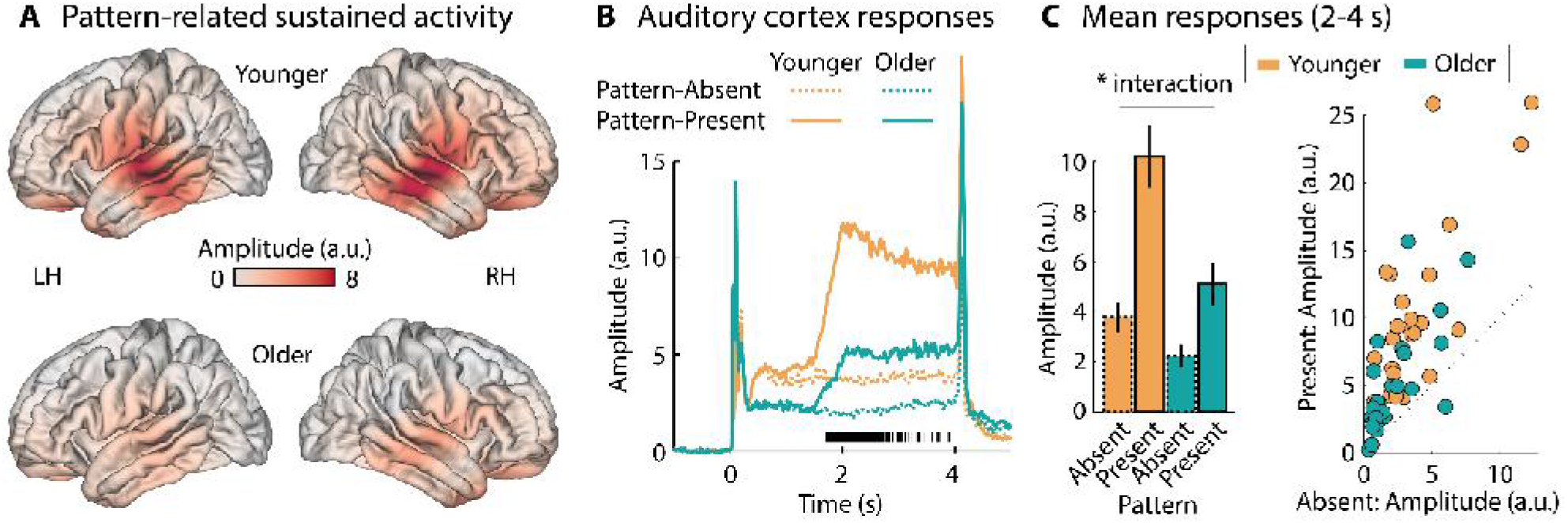
Source localization of pattern-evoked sustained activity. A: Source localization of pattern-related sustained activity (difference between Pattern-Present and Pattern-Absent conditions). B: Response time courses from auditory cortex. The black markings below the time courses indicate the time points at which the Condition × Age Group interaction was significant (FDR-thresholded). C: Mean auditory cortex responses in the 2-4 s time window. Bar graphs reflect the mean across participants. Error bars are the standard error of the mean. Data points for each participant are shown on the right. Data points above the diagonal (dashed line) reflect a larger response for the Pattern-Present compared to the Pattern-Absent condition.

## Discussion

The current magnetoencephalography study investigated age-related differences in auditory cortical responsivity to sound onsets and to the presence of a pattern in sounds. We showed that older adults elicit larger responses in auditory cortex to sound onsets compared to younger adults. This response enhancement indicates that auditory cortex of older adults is hyperresponsive to sound. Despite this age-related hyperresponsiveness, sustained neural activity in auditory cortex to sound patterns was diminished in older compared to younger people. Our results suggest that neural responses in auditory cortex are fundamentally altered in older adults such that cortical activity overrepresents sound onsets, whereas it underrepresents temporally coherent structure in sounds.

### Hyperresponsiveness of auditory cortex in older adults

We demonstrated that neural responses in the M50 and M100 time window following sound onset are enhanced in older compared to younger adults (Figure 3A-C). We localized the M50 and M100 responses to auditory cortex (Figure 3D/E; consistent with previous work Pantev et al., 1988; Maess et al., 2007; Okamoto and Kakigi, 2014; Herrmann et al., 2018), suggesting that auditory cortex in older adults is hyperresponsive. This is in line with a growing literature showing that neural responses to sound onsets are enhanced in older compared to younger adults (Ross and Tremblay, 2009; Sörös et al., 2009; Lister et al., 2011; Alain et al., 2012; Bidelman et al., 2014; Herrmann et al., 2016; Herrmann and Johnsrude, 2018a). Similar observations have been made for aged monkeys (Juarez-Salinas et al., 2010; Recanzone, 2018) and aged rodents (Hughes et al., 2010), as well as for non-human mammals whose auditory periphery was damaged through high-intensity sound exposure (Popelár et al., 1987; Syka et al., 1994; Schormans et al., 2019) or ototoxic drugs (Qiu et al., 2000; for detailed reviews see Auerbach et al., 2014; Zhao et al., 2016; Salvi et al., 2017; Herrmann and Butler, 2021).

Hyperresponsiveness to sound is thought to result from hyperexcitable neural circuits due to a loss of inhibition in the auditory system following peripheral decline (Caspary et al., 2008; Takesian et al., 2012). The functional role of the loss of inhibition and hyperexcitability is still debated (Zhao et al., 2016; Asokan et al., 2018; Herrmann and Butler, 2021), but likely includes homeostatic processes to regulate excitation (Caspary et al., 2008; Zhao et al., 2016) and a state of increased plasticity that enables cortical reorganization (Cisneros-Franco et al., 2018; Cisneros-Franco and de Villers-Sidani, 2019). A balanced level of excitation and inhibition is crucial for neural function (Wehr and Zador, 2003; Silver, 2010; Isaacson and Scanziani, 2011), and the fact that we observed hyperresponsiveness to sound in older compared to younger adults suggests that neural function of auditory cortex was altered in our sample of older individuals. Hyperresponsivity to sharp attacks in sound may underlie increased distractibility by irrelevant sounds in older compared to younger adults (Parmentier and Andrés, 2010) and difficulties comprehending speech in the presence of an interfering, modulated background masker (Millman et al., 2017; Goossens et al., 2018).

### Pattern-related activity is reduced in older compared to younger adults

In order to investigate whether neural sensitivity to a pattern in sounds differs between younger and older adults, we presented sounds that either contained a pattern (made of a sequence of a repeated set of pure tones at different frequencies) or did not contain a pattern (made of a sequence of tones at pseudo-randomly selected frequencies; Figure 2). For both younger and older adults, we observed that sustained neural activity increased after the onset of a sound pattern relative to sounds without a pattern. Previous work in younger adults has revealed similar increases in sustained activity for different types of patterns, including tone sequences such as those we have utilized here (Gutschalk et al., 2002; Ross et al., 2002; Keceli et al., 2012; Barascud et al., 2016; Sohoglu and Chait, 2016; Teki et al., 2016; Southwell et al., 2017; Herrmann and Johnsrude, 2018b; Southwell and Chait, 2018; Herrmann et al., 2019; Herrmann et al., 2021).

We showed that sustained neural activity to a pattern in sounds is reduced in older compared to younger adults. Hence, although neural responses to the onset of sound was enhanced in older adults, neural sensitivity to a pattern in sounds was reduced. Diminished sustained activity for older compared to younger adults is consistent with previous indications of an age-related reduction in sustained activity for short (<1 s) pure tones (Pfefferbaum et al., 1979), amplitude modulations (Herrmann et al., 2019), and repeated patterns in tone sequences (Al Jaja et al., 2020). However, low statistical reliability and differences in stimulus parameters between age groups did not allow drawing firm conclusions from the latter two studies. Our results demonstrate clearly that pattern-related sustained activity indeed is reduced in older adults.

Sensitivity to sound patterns is crucial for a variety of auditory functions, enabling a listener to segregate concurrent sound streams (Schröger, 2005, 2007; Snyder and Alain, 2007; Winkler et al., 2009; Bendixen, 2014) and recognize and predict relevant sounds (Jones and Boltz, 1989; Nobre et al., 2007; Henry and Herrmann, 2014; Nobre and van Ede, 2018). By demonstrating a correlation between perceptual sensitivity to sound patterns and speech comprehension abilities (Holmes and Griffiths, 2019) and common substrates in auditory cortex (Holmes et al., 2021), previous work further indicates a functional relation or common underlying mechanism between the processing of regularities in sounds and speech comprehension. A reduction in sustained activity may thus indicate that sound patterns are processed less well in neural circuits in older compared to younger adults, which may, in part, explain the challenges older adults experience comprehending speech in the presence of background sound.

Participants in the current study were presented with sound sequences while they watched a muted, subtitled movie of their choice. Participants’ attention was thus directed away from the sounds, although the degree of attentional focus was not experimentally constrained in the current study. Previous work indicates that regularity-related sustained activity can be increased if participants perform a difficult sound-related task relative to a difficult visual task (Herrmann and Johnsrude, 2018b). Larger regularity-related sustained activity in younger compared to older adults could thus be, in part, the result of younger adults attending more to the sounds than older adults. However, younger and older adults typically enjoy watching a movie in such experiments, where they are not required to perform a sound-related task. Moreover, larger responses in older compared to younger adults to the sound onset may indicate greater attentional capture by sounds for older adults (see also Parmentier and Andrés, 2010; Weeks and Hasher, 2014), but regularity-related sustained activity was decreased for them. Differences in the degree of attention to sounds between age groups are thus unlikely to explain the observed differences in regularity-related sustained activity.

The current source localizations suggest that auditory cortex is the main source underlying pattern-related sustained activity in both younger and older adults (Figure 5A). Previous work in younger individuals also indicated that auditory cortex underlies sustained neural activity (Hari et al., 1980; Pantev et al., 1994; Pantev et al., 1996; Gutschalk et al., 2002; Ross et al., 2002; Gutschalk et al., 2004; Gutschalk et al., 2007; Okamoto et al., 2011; Keceli et al., 2012; Barascud et al., 2016; Teki et al., 2016), but that brain regions in frontal cortex, parietal cortex, and hippocampus may additionally contribute (Tiitinen et al., 2012; Barascud et al., 2016; Teki et al., 2016). However, in the latter work, statistical difference maps were calculated and used to identify neural sources. Statistical difference maps may also capture effects related to activity spread due to volume conduction and may thus not reflect activity originating from these higher-level brain regions (e.g., auditory responses to sound onset were spread to parietal cortex in Teki et al., 2016, suggesting that spread may also affect their sustained activity in parietal cortex related to sound patterns).

We further showed that sustained activity in auditory cortex to sounds that did not contain a pattern was also reduced in older compared to younger adults (Figure 5B/C). Sounds without a pattern were made of a sequence of pure tones whose frequency changed randomly for each tone. Such tone sequences are perceptually more structured than noise and the auditory system may treat them as a pattern of low saliency. This is consistent with the observation of reduced sustained activity to short pure tones in older compared to younger adults (Pfefferbaum et al., 1979). Our data thus indicate that the sensitivity of the aged auditory cortex is reduced for sounds containing a pattern (here repetition of a set of tones at different frequencies) as well as for sequences with random tone frequencies.

It is clear from previous work that temporally regular – and thus predictable – structure in sounds that forms a pattern elicits sustained neural activity (Gutschalk et al., 2002; Barascud et al., 2016; Herrmann and Johnsrude, 2018b). However, additional work suggests that the magnitude of pattern-related sustained activity is related to the degree of novelty or predictability of a pattern, such that sustained activity decreases when a pattern is frequently, compared to infrequently, heard (Gutschalk et al., 2007; Herrmann et al., 2021). A reduction in sustained activity in older adults may thus result from reduced processing of the pattern as well as from reduced novelty of the pattern, but further behavioral research is needed to investigate the perceptual consequences of the altered cortical sensitivity observed here.

## Conclusions

In the current study, we recorded magnetoencephalography to characterize differences between younger and older adults in the processing of a pattern in sounds. We presented continuous tone sequences that either contained a pattern (made of a repeated set of tones at different frequencies) or did not contain a pattern (random tone frequencies). We showed that auditory cortex in older adults is hyperresponsive to sound onsets, but that sustained neural activity in auditory cortex, indexing the processing of sound patterns, is reduced. Hence, neural populations in auditory cortex fundamentally differ between younger and older individuals in their sensitivity to sound features, hyperresponding to sound onsets, while underresponding to patterns in sounds. This may help to explain some age-related changes in hearing such as increased sensitivity to distracting sounds and difficulties tracking speech in the presence of other sound.

## Acknowledgements

Research was supported by a Canadian Institutes of Health Research (MOP133450) grant to ISJ. BH was supported by a BrainsCAN postdoctoral fellowship (Canada First Research Excellence Fund; CFREF) and the Canada Research Chair program. We thank the Max Planck Institute for Human Cognitive and Brain Sciences for the opportunity to record the data. We thank Yvonne Wolff-Rosier for help during data acquisition.

## Author contributions

BH conceptualized and designed the study, recorded data, analyzed the data, interpreted the results, and wrote the manuscript. BM analyzed the data, interpreted the results, and edited the manuscript. ISJ conceptualized and designed the study, interpreted the results, and edited the manuscript.

## Footnotes

1 The dB values are derived from MATLAB. More negative values reflect softer sound intensities. These dB values can be interpreted relative to each other, whereas the absolute magnitude is related to hardware and software conditions, such as sound card, transducers, and MATLAB internal settings.

